# Endogenous retrovirus-driven *Pcgf5* plays critical roles in zygotic genome activation and noncanonical imprinting

**DOI:** 10.1101/2025.01.24.632300

**Authors:** Satoshi Mashiko, Takuto Yamamoto, Naojiro Minami, Shuntaro Ikeda, Shinnosuke Honda

## Abstract

MERVL (murine endogenous retrovirus with leucine tRNA primer) is expressed during zygotic genome activation (ZGA) in mammalian embryos. Here, we show that the Polycomb group ring finger 5 (*Pcgf5*), a key component of Polycomb repressive complex 1 (PRC1), forms a chimeric transcript with MT2C_Mm, one of the long terminal repeat sequences of MERVL. Knockdown of *Pcgf5* reduced developmental rates and decreased H3K27me3 and H2AK119ub1 modification during embryogenesis. In addition, not only genes expressed during ZGA but also imprinting genes were upregulated in *Pcgf5* knockdown embryos. Moreover, *Pcgf5* was involved in the addition of the H3K27me3 modification to the maternal *Xist* region. This is the first report of a MERVL-regulated transcript regulating *Xist* expression in mouse preimplantation embryos. Our results suggest that analysis of chimeric transcripts with MERVL will provide insight into the relationship between ZGA and noncanonical imprinting.

## Introduction

In early mammalian embryogenesis, the development of embryos after fertilization is supported by maternal factors stored in the oocyte that trigger zygotic genome activation (ZGA).^1^ ZGA has two waves: minor and major. The former occurs during the S phase of the one-cell stage while the latter occurs during the late two-cell stage.^2,3^ Before and after ZGA, the epigenome of the embryos changes dramatically.^4,5^ Epigenetic modifications such as DNA methylation and histone modifications are how gene expression is regulated and maintained without dependence on a genome sequence and are considered to be important for reprogramming and cell differentiation during embryo development.^6^ After the major ZGA, cells differentiate into the inner cell mass and the trophectoderm as development proceeds to the blastocyst stage.

The long terminal repeat (LTR) of endogenous retrovirus (ERV) expresses chimeric mRNAs involving adjacent genes, and ERVs are predicted to regulate the expression of other genes.^7^ Murine endogenous retrovirus with leucine tRNA primer (MERVL) is an ERV expressed during ZGA. Although transcription of the MERVL retrotransposon is required for preimplantation embryo development, the function of the vast majority of MERVL chimeric mRNAs remains unclear.^8^ The mammalian genome is rich in ERVs,^9^ suggesting that ERVs were involved in the evolution of the placenta in mammals.

Because MERVL is reported to be one of the earliest genomic sequences transcribed in the preimplantation embryo,^10^ we hypothesized that the MERVL chimeric transcript plays a major role in preimplantation embryo development. In our laboratory, reanalysis of previously reported RNA-seq data revealed that expression of protein arginine methyltransferase 6 (*Prmt6*) is activated by the MERVL promoter sequence in the preimplantation embryo and generates chimeric and canonical transcripts with distinct functions.^11^ In the same analysis, Polycomb group ring finger 5 (*Pcgf5*) was found to be activated by MERVL in preimplantation embryos. *Pcgf5* forms one of the four subtypes of Polycomb repressive complex 1 (PRC1). The essential core of PRC1 consists of an E3 ubiquitin ligase, either RING1A or RING1B. There are six different forms of PRC1 (PRC1.1–PRC1.6), each of which contains six different Polycomb group ring finger (PCGF) proteins (PCGF1–PCGF6) that are essential for PRC1 function.^12^ Among these, variant PRC1 (vPRC1) proteins (PRC1.1, PRC1.3, PRC1.5, and PRC1.6) have high catalytic activity for the monoubiquitination of histone H2A at lysine 119 (H2AK119ub1).^13,14^

In addition to DNA methylation, another modification that is passed down from gametes to the next generation is trimethylation of histone H3 at the 27th lysine (H3K27me3), a phenomenon called noncanonical imprinting.^15–17^ Moreover, at specific genomic regions, H3K27me3 in oocytes is maintained in maternal chromosomes after fertilization, represses gene expression, and contributes to proper placental formation.^15,17^ Several paternally expressed genes (PEGs), such as *Sfmbt2* and *Slc38a4*, have been identified as noncanonical imprinting genes.^15,18–20^ In mice, maternal H3K27me3 mediates not only autosomal maternal allele-specific gene silencing but also plays an important role in X chromosome inactivation (XCI) through repression of maternal *Xist.*^16,21^ Recently, analysis of *Pcgf1* and *Pcgf6* knockout (KO) mice revealed that H2AK119ub1, which is modified in oocytes via PRC1 containing PCGF1/6, colocalizes with the H3K27me3 modification in mouse embryos.^22^ While the maternal H3K27me3 domains are maintained throughout preimplantation development, H2AK119ub1 at these loci undergoes transient loss at the two-cell stage and is then re-established allelically at later stages.^23^ In addition, *Pcgf3/5* KO embryonic stem cells fail to accumulate H3K27me3 on the inactive X chromosome.^24,25^ However, the function of *Pcgf5* in the preimplantation embryo and the significance of the generation of chimeric *Pcgf5* transcripts remain unclear.

Here, we show that *Pcgf5* and chimeric *Pcgf5* expressed may be involved in ZGA gene regulation and XCI, as well as histone modification. MT2C_Mm is one of the LTRs of MERVL and RNA-seq data provided by Macfarlan et al. showed that MT2C_Mm-driven *Pcgf5* (*Pcgf5^MT2C_Mm^*) is highly expressed during ZGA.^26^ Most of the *Pcgf5* expressed at the two-cell stage is regulated by MERVL sequences, and knockdown (KD) with siRNA targeting the *Pcgf5* common exon resulted in lower developmental rates from the four-cell stage onward. Furthermore, *Pcgf5* siRNA KD embryos showed abnormal expression of ZGA genes and *Xist* expression and fewer H2AK119ub1 and H3K27me3 modifications during embryogenesis. In addition, allele-specific ChIP-seq analysis confirmed decreased H3K27me3 modification in the maternal *Xist* allele. KO mice lacking the MERVL LTR sequence for transcribing chimeric *Pcgf5* were fertile, but the developmental competence of the KO embryos was reduced. Furthermore, severe KD of *Pcgf5* using antisense oligonucleotides (ASOs) completely arrested embryogenesis in two- to four-cell embryos. Taken together, these results suggest that ERVs play a major role in appropriate embryogenesis in terms of genomic imprinting and ZGA.

## Results

### Expression levels of *Pcgf5* and chimeric *Pcgf5* in preimplantation embryos

Analysis of public RNA-seq data^27^ showed that transcription of *Pcgf5^MT2C_Mm^* had begun at the late two-cell stage (Fig. S1). While four canonical *Pcgf5* variants (*Pcgf5^CAN^*) (NM_001368690.1, NM_029508.4, NM_001360691.1, and XM_036161722.1; hereafter, NM90, NM08, NM91, and XM22, respectively) are reported in the NCBI database (https://www.ncbi.nlm.nih.gov/), RepeatMasker (https://www.repeatmasker.org/) annotation data for the mouse genome (mm10) showed that MT2C_Mm was localized between the first exons of NM08 to NM91 (Fig. 1A). To determine the expression profiles of *Pcgf5^CAN^* and *Pcgf5^MT2C_Mm^*, we performed quantitative reverse-transcription PCR (RT-qPCR) at each developmental stage (Fig. 2A). The results showed that total *Pcgf5* (both *Pcgf5^CAN^* and *Pcgf5^MT2C_Mm^*) and *Pcgf5^MT2C_Mm^* expression increased in the late two-cell stage and gradually decreased in the blastocyst stage (Fig. 1B). To compare the copy number of each transcript, absolute quantification was performed using late two-cell embryos and blastocysts. In late two-cell embryos, both total *Pcgf5* and *Pcgf5^MT2C_Mm^* copy numbers were almost the same (Fig. 1C). However, in blastocysts, the copy number of *Pcgf5^MT2C_Mm^* was significantly lower than that of total *Pcgf5* transcripts (Fig. 1C).

**Fig. 1.**
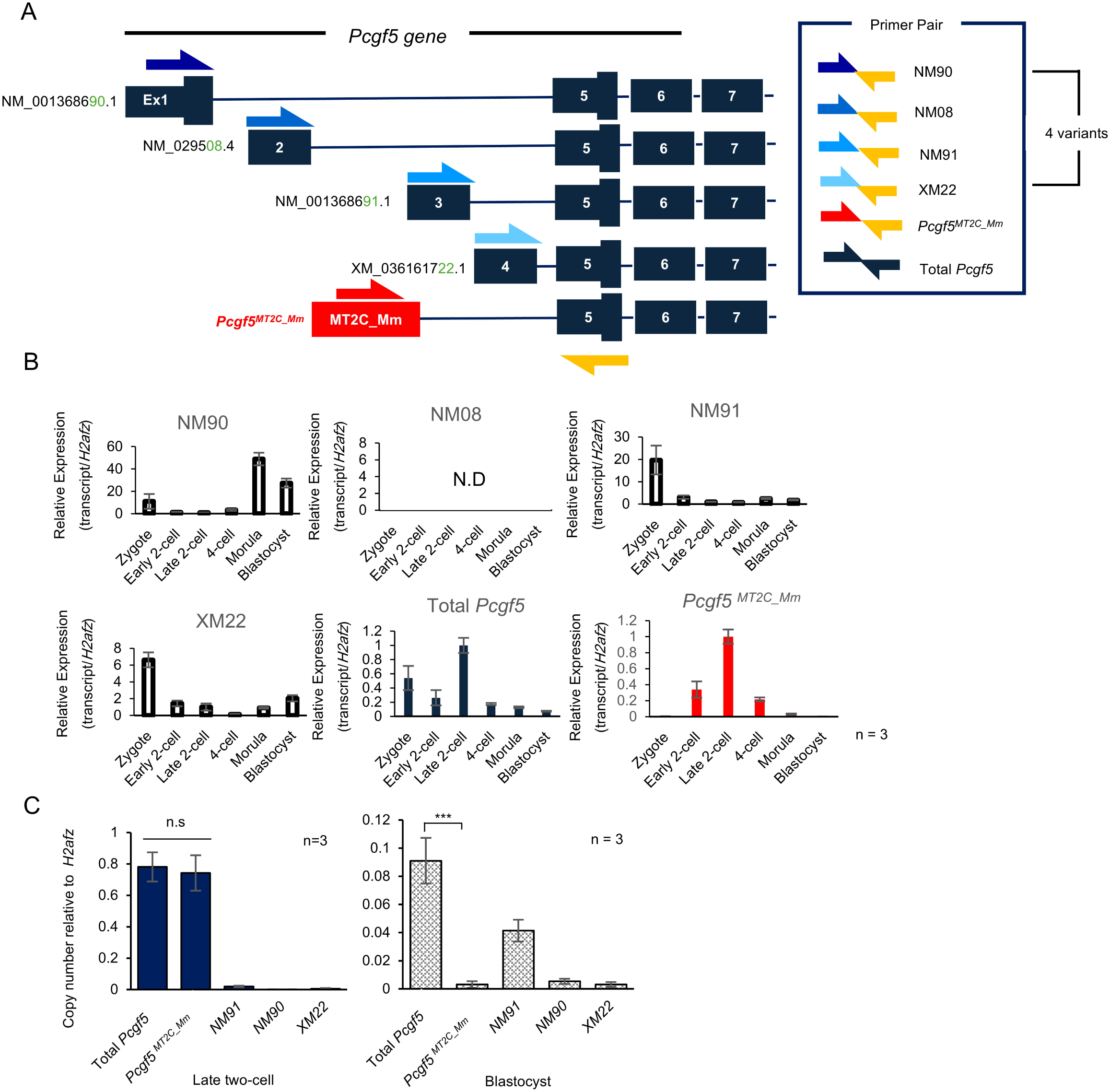
Gene structure and transcriptional profile of *Pcgf5* in mice. (A) Schematic diagram showing the positions of MT2C_Mm and *Pcgf5* exons in the mouse genome. *Pcgf5* has four canonical variants (NM_001368690.1, NM_029508.4, NM_001360691.1, and XM_036161722.1) and the primer pairs for detecting these mRNAs were named NM90, NM08, NM91, and XM22, respectively. Similarly, the primers used to detect chimeric *Pcgf5* and total *Pcgf5* mRNA expression were named *Pcgf5^MT2C_Mm^* and total *Pcgf5*, respectively. (B) RT-qPCR analysis of each *Pcgf5* variant, total *Pcgf5*, and *Pcgf5^MT2C_Mm^* transcripts during mouse preimplantation development. mRNA expression levels were normalized to *H2afz* as an internal control. Data are expressed as mean ± s.e.m. (n = 3). N.D., not detected. (C) Comparison of the expression levels of total *Pcgf5*, each *Pcgf5* variant, and *Pcgf5^MT2C_Mm^* in late two-cell embryos and blastocysts based on absolute quantification by RT-qPCR. Each graph shows the expression levels of each transcript in late two-cell embryos and blastocysts. Copy numbers were normalized to *H2afz* copy numbers. Data are expressed as mean ± s.e.m. (n = 5). Two-tailed Student’s t-test: ***P < 0.001; n.s., not significant.

**Fig. 2.**
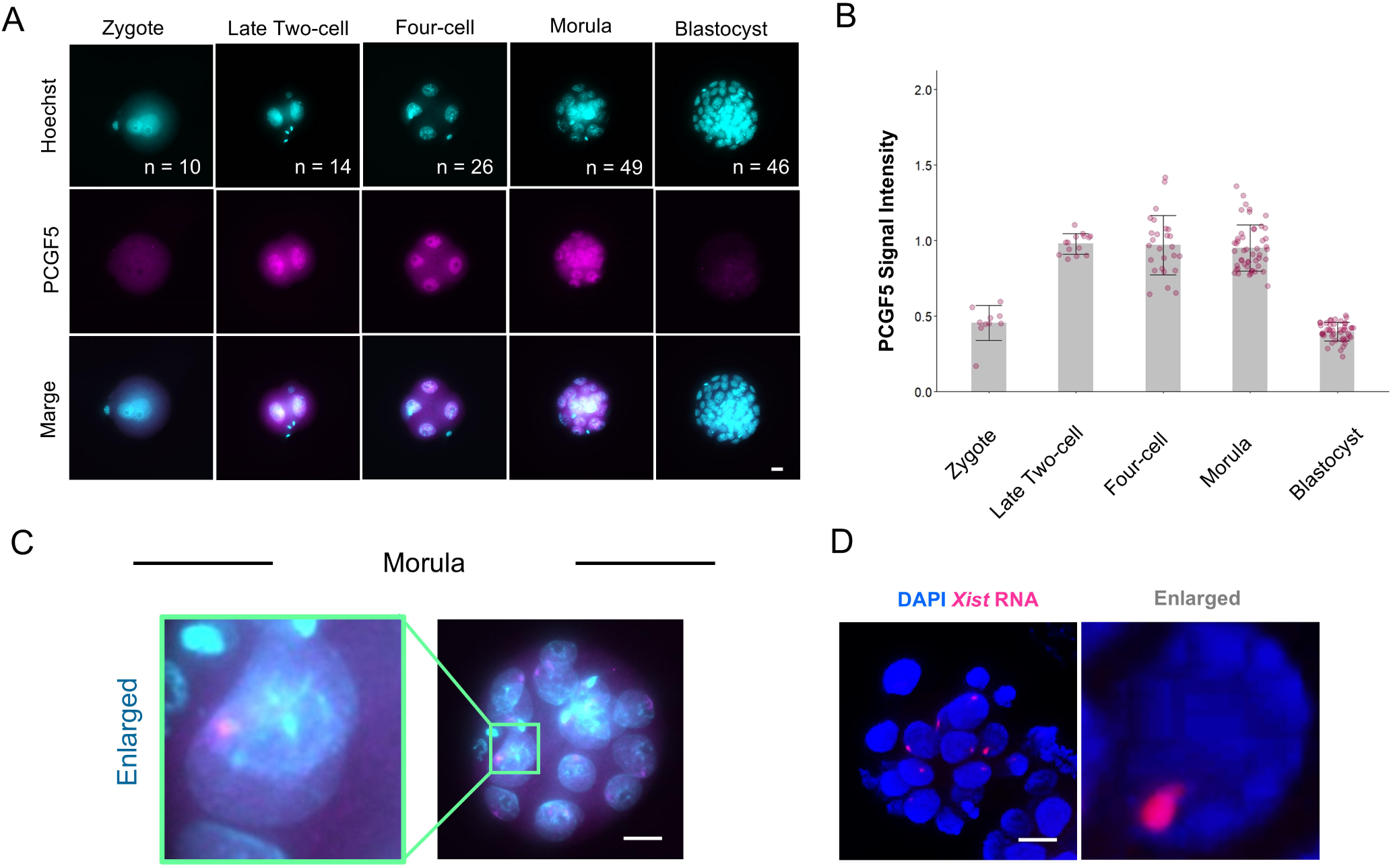
Expression of PCGF5 protein. (A) Representative images of PCGF5 immunostaining analysis at each stage of mouse preimplantation embryos. The numbers of embryos examined (n) are indicated. Scale bar, 20 μm. (B) Quantification of PCGF5 immunostaining analysis at each stage of embryos. The numbers of embryos examined (n) are indicated in Fig. 2A. The average signal intensity of late two-cell embryos was set as 1.0 in each stage. Bars show means with standard errors of the mean (s.e.m.). (C) Only one PCGF5 spot was identified per blastomere in some morulae. The arrowhead indicates a significant PCGF5 spot. The left panel shows an enlarged view of a clear PCGF5 spot. Scale bar, 20 μm. (D) Only one *Xist* spot was identified per blastomere in some morulae, similar to PCGF5 spots. The right panel shows an enlarged view of a clear *Xist* spot. Scale bar, 20 μm.

### Dynamics of PCGF5 protein expression in preimplantation embryos

Immunofluorescence revealed that PCGF5 expression peaked in the late two-cell stage and was maintained until the morula stage. In the blastocyst stage, PCGF5 expression decreased to the same level as in the zygote (Fig. 2A, B). In addition, interestingly, one PCGF5 spot was observed per blastomere only in about half of morulae, and *Xist* FISH (fluorescence in situ hybridization) analysis similarly revealed a *Xist* spot in about half of morulae (Fig. 2C,D).

### *Pcgf5* knockdown experiments using siRNA

Electroporation with siRNAs targeting the *Pcgf5* exon sequence (si*Pcgf5^CAN^*-1 or *siPcgf5^CAN^*-2) resulted in a significant decrease in the transcript levels of both total *Pcgf5* and *Pcgf5^MT2C_Mm^* transcripts at the late two-cell stage (Fig. 3A). The developmental competence of *Pcgf5* siRNA KD embryos decreased significantly after the four-cell stage compared with embryos electroporated with siControl (CTR) (Fig. 3B, C). PCGF5 expression in KD embryos was significantly decreased at the late two-cell stage compared with CTR embryos, but the PCGF5 signal recovered after the four-cell embryo stage (Fig. 3D, E).

**Fig. 3.**
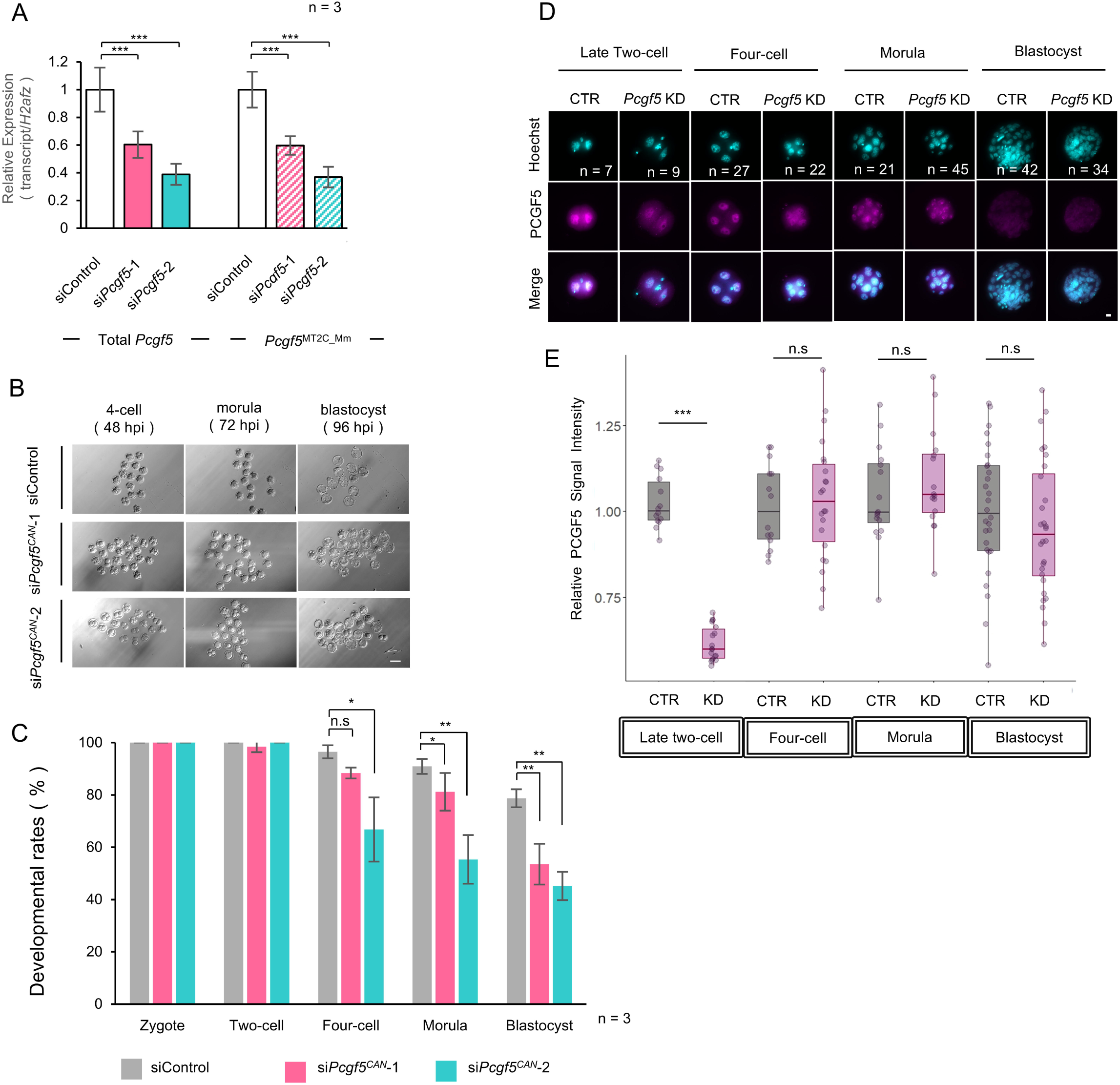
A decrease in *Pcgf5* causes a reduced developmental rate. (A) RT-qPCR analysis of total *Pcgf5* and *Pcgf5^MT2C_Mm^*mRNA levels in late two-cell embryos electroporated with si*Pcgf5^CAN^*-1, si*Pcgf5^CAN^*-2, or siControl at 3 hpi. The amount of transcripts in siControl-electroporated embryos was defined as 1, and the expression levels were normalized to *H2afz* as an internal control. Data are expressed as mean ± s.e.m. (n = 3). Dunnett’s test: ***P < 0.001. (B) Representative images of preimplantation embryos that had been electroporated with si*Pcgf5^CAN^*-1, si*Pcgf5^CAN^*-2, or siControl at 3 hpi at the indicated times. Scale bar, 100 μm. (C) Preimplantation development rates of embryos electroporated with si*Pcgf5^CAN^*-1, si*Pcgf5^CAN^*-2, or siControl. The embryos that reached the two-cell, four-cell, morula, and blastocyst stages were counted at 24, 48, 72, and 96 hpi, respectively. The numbers of zygotes were set as 100%. The numbers of embryos examined were 51 (si*Pcgf5^CAN^*-1), 60 (si*Pcgf5^CAN^*-2), and 48 (siControl) from three biologically independent experiments. Data are expressed as mean ± s.e.m. (n = 3). Dunnett’s test: *P < 0.05, **P < 0.01; n.s., not significant. (D) Representative images of PCGF5 immunostaining analysis in siControl-electroporated (CTR) and *Pcgf5* knockdown (KD) embryos. Late two-cell embryos, four-cell embryos, morulae, and blastocysts were collected at 36, 48, 72, and 96 hpi, respectively. The numbers of embryos examined (n) are indicated. Scale bar, 20 μm. (E) Quantification of PCGF5 immunostaining analysis in CTR and *Pcgf5* KD embryos. The numbers of embryos examined (n) are indicated in Fig. 3D. The average signal intensity of CTR was set as 1.0 in each stage. In box plots, the center lines correspond to the median, the lower and upper edges of boxes to the first and third quartiles, and the whiskers to the minimum and maximum values. Two-tailed Student’s t-test: ***P < 0.001; n.s., not significant.

### Functional analysis of *Pcgf5* during ZGA

RT-qPCR was performed on the expression of four major transcripts during ZGA— *MERVL*, *Zscan4*, *Zfp352*, and *Msfd7c*—using samples from CTR and siRNA KD embryos at the late two-cell and the four-cell stages. There was no significant change in the expression of these four genes at the late two-cell stage (Fig. 4A). In contrast, *Zscan4* and *Zfp352* were significantly upregulated in siRNA KD embryos at the four-cell stage (Fig. 4B).

**Fig. 4.**
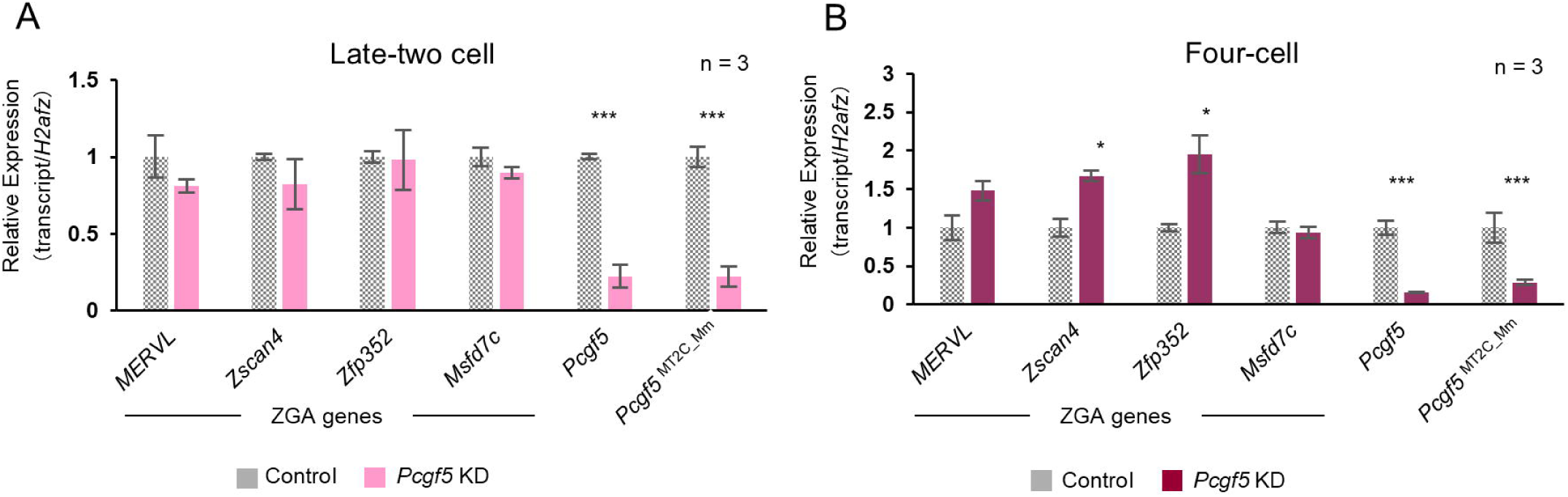
Genes expressed during ZGA as measured by RT-qPCR in si*Pcgf5* KD embryos. (A) Genes expressed during ZGA as measured by RT-qPCR in siControl and si*Pcgf5* KD late two-cell embryos at 36 hpi. There were no significant changes in the expression of four major ZGA genes. (B) Genes expressed during ZGA as measured by RT-qPCR in siControl and si*Pcgf5* KD four-cell embryos at 48 hpi. *Zscan4* and *Zfp352* were significantly upregulated in si*Pcgf5* KD embryos. Two-tailed Student’s t-test: *P < 0.05, ***P < 0.001.

### Functional analysis of *Pcgf5* in genomic imprinting

Co-immunofluorescence of H2AK119ub1 and H3K27me3 showed that both signals were significantly lower in siRNA KD embryos at the late two-cell and the four-cell stage compared with CTR embryos (Fig. 5A, B). RT-qPCR of imprinted genes in morulae showed that *Pcgf5* siRNA KD embryos had significantly higher expression of *Sfmbt2* and *Xist* among the 18 genes examined (Fig. 5C). Chromatin immunoprecipitation sequencing (ChIP-seq) revealed that the peak of the H3K27me3 modification in *Xist* was lower in *Pcgf5* siRNA KD morulae than in CTR and the maternal allele of *Xist* had decreased H3K27me3 modification (Fig. 5D).

**Fig. 5.**
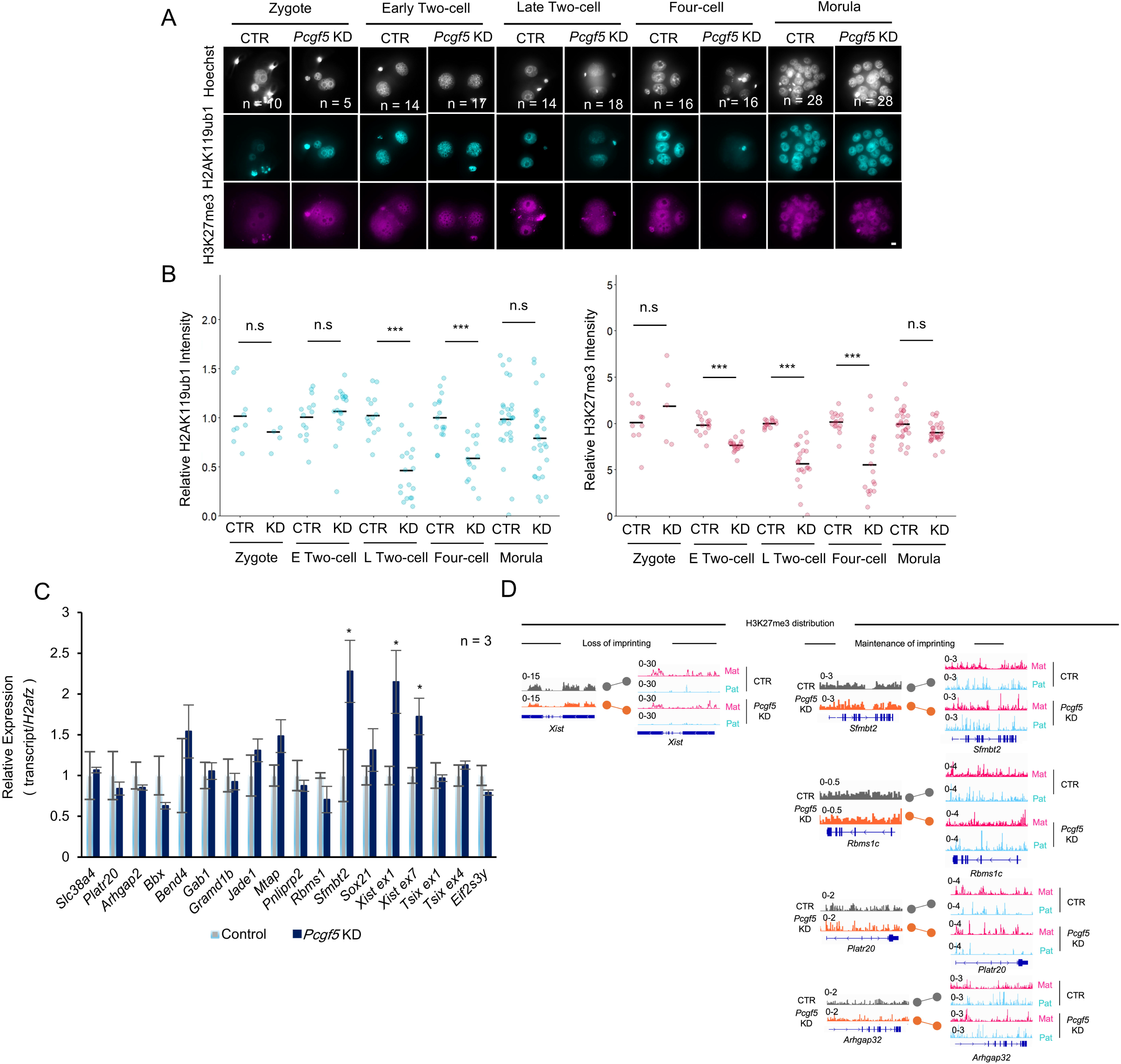
Analysis of histone modifications in si*Pcgf5* KD embryos. (A) Representative images of H2AK119ub1 and H3K27me3 immunostaining analysis in siCTR and si*Pcgf5* KD embryos. Zygotes, early and late two-cell embryos, four-cell embryos, and morulae were collected at 6, 24, 36, 48, and 72 hpi, respectively. The numbers of embryos examined (n) are indicated. Scale bar, 20 μm. (B) Quantification of H2AK119ub1 and H3K27me3 immunostaining analysis in siCTR and si*Pcgf5* KD embryos. The numbers of embryos examined (n) are indicated in Fig. 5B. The average signal intensity of CTR was set as 1.0 in each stage. Bars overlaid on the plots indicate the mean. Two-tailed Student’s t-test: ***P < 0.001; n.s., not significant. (C) Maternal noncanonical imprinting genes as measured by RT-qPCR in siCTR and si*Pcgf5* KD morulae at 72 hpi. *Xist* ex1 or ex7 refers to the exon 1 or exon 7 region of the *Xist* gene, and the same applies to *Tsix*. Bars show means with standard error of the mean (s.e.m.). Biological replicates (n) are indicated. Two-tailed Student’s t-test: *P < 0.05. (D) Genome browser views of H3K27me3 distributions in siCTR and si*Pcgf5* KD morulae. Mat, maternal allele; Pat, paternal allele. The scales of the tracks are indicated on the left of each track.

### Analysis of *Pcgf5* KO mice and chimeric *Pcgf5* KO mice

To verify the role of chimeric *Pcgf5* in embryogenesis, two KO mouse strains were generated: those lacking MERVL LTR to transcribe chimeric *Pcgf5* (*Pcgf5^ΔMT2C_Mm/+^*) and those with deletion of exon 5 of *Pcgf5* (*Pcgf5^ΔEx^*^5^*^/+^*) (Fig. 6A). The *Pcgf5^ΔEx^*^5^*^/+^* mice showed no difference in litter size in heterozygous matings but were unable to produce homozygous KO mice (Fig. 6B, C). In contrast, *Pcgf5^ΔMT2C_Mm/+^* mice exhibited no change in litter size or in their potential to produce homozygous KO mice (*Pcgf5^ΔMT2C_Mm/ΔMT2C_Mm^*) (Fig. 6B, C). However, *Pcgf5^ΔMT2C_Mm/ΔMT2C_Mm^* mice were significantly less likely to produce females (Fig. 6D). This phenomenon was also observed in *Pcgf5^ΔEx^*^5^*^/+^* mice, which were similarly less likely to produce female (Fig. 6E). Subsequently, the developmental rate of *Pcgf5^ΔMT2C_Mm/ΔMT2C_Mm^*embryos was found to be mildly decreased at the blastocyst stage (Fig. 6F, G). Furthermore, immunofluorescence of PCGF5, H2AK119ub1, and H3K27me3 protein showed significant reduced fluorescence intensity in both two- and four-cell embryos, as well as in *Pcgf5* KD embryos (Fig. 6H, I). To determine why the *Pcgf5^ΔMT2C_Mm/ΔMT2C_Mm^* mice had a milder phenotype than the *Pcgf5^ΔEx^*^5^*^/ΔEx^*^5^ mice, we performed in vitro fertilization using *Pcgf5^ΔMT2C_Mm/ΔMT2C_Mm^* mice and conducted RT-qPCR at the late two-cell, four-cell, and morula stages. Although there were no significant changes in the expression of imprinted genes, several genes expressed during ZGA (i.e., *Dub1* and *Tcstv3*) were significantly upregulated in *Pcgf5^ΔMT2C_Mm/ΔMT2C_Mm^* embryos (Fig. S2). In the case of the expression of *Pcgf5* variants, including *Pcgf5^MT2C_Mm^*, the halved expression of total *Pcgf5* was rescued at the late two-cell stage, even though the expression of total *Pcgf5* was mostly dependent on *Pcgf5^MT2C_Mm^* expression at this stage (Fig. 6J). This finding was supported by the approximately 3-fold upregulation of NM91, one of the variants of *Pcgf5^CAN^*, suggesting that *Pcgf5^CAN^* can rescue total *Pcgf5* expression even when MT2C_Mm is completely deleted (Fig. 6H).

**Fig. 6.**
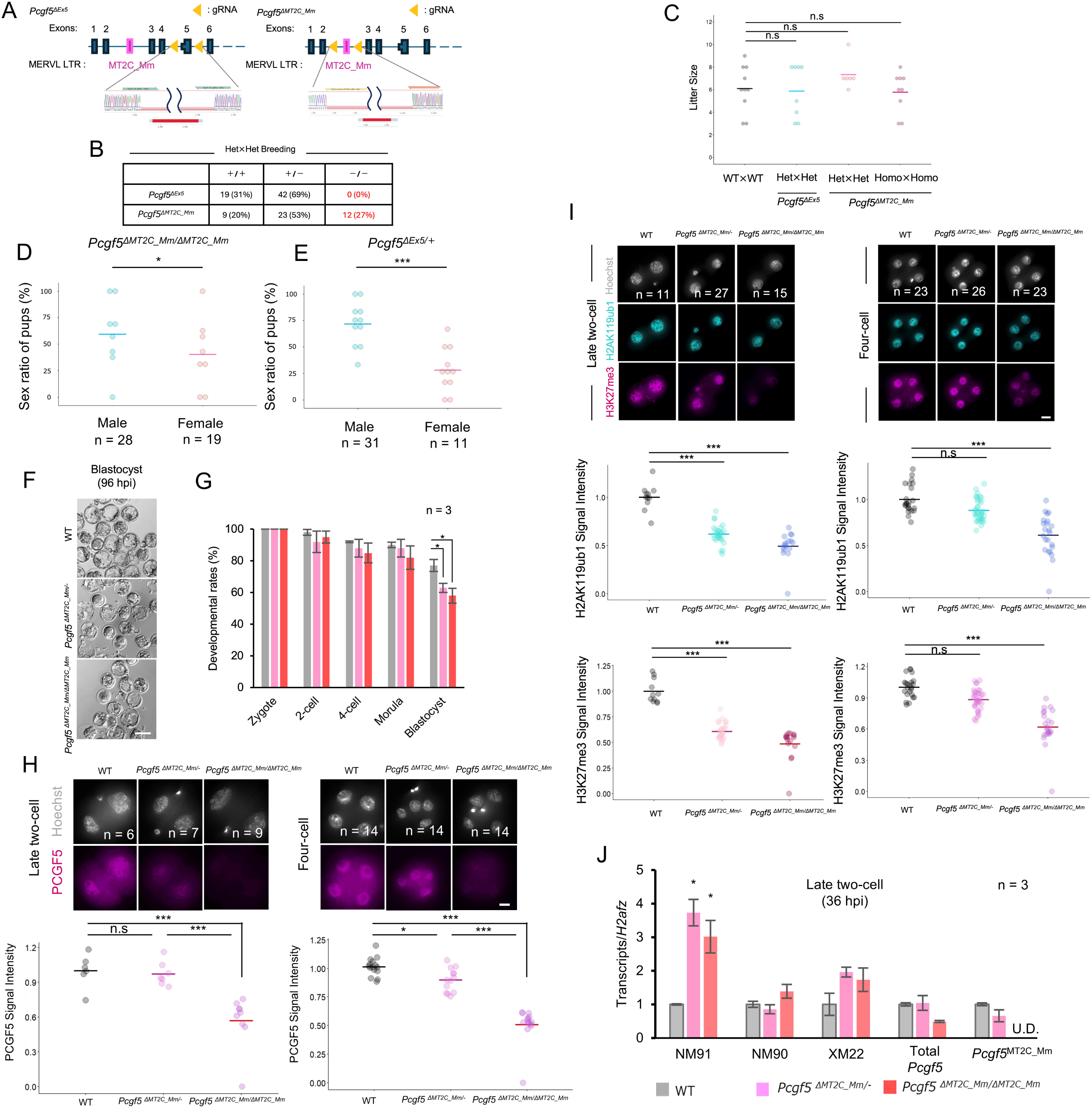
Analysis of *Pcgf5* KO and MT2C_Mm KO mice. (A) Strategy for CRISPR-Cas9-mediated mutagenesis of *Pcgf5* exon 5 and MT2C_Mm loci. The positions of guide RNAs relative to *Pcgf5* exon 5 and MT2C_Mm are shown. An alignment of the deletions of the strains used in this study are shown below. (B) Mating results between the heterozygous KO mice obtained. +/+, +/−, and −/− indicate that the genotyped pups were wild-type, heterozygous, and homozygous KO (red font). Numbers in the box indicate the number of pups. (C) Litter size of knockout mice. A single dot indicates the number of pups. Gray indicates results for wild-type (WT), blue for *Pcgf5* exon 5 KO mice and red for MT2C_Mm KO mice. One dot indicates the number of litters from one mother. Bars overlaid on the plots indicate the mean. Dunnett’s test: n.s., not significant. (D) Sex ratio of pups obtained from crosses between *Pcgf5^ΔMT2C_Mm/ΔMT2C_Mm^* mice. The numbers of pups examined (n) are indicated in Fig. 6C. (E) Sex ratio of pups obtained from crosses between *Pcgf5^ΔEx^*^5^*^/+^* mice. The numbers of pups examined (n) are indicated in Fig. 6C. (F) Brightfield photograph of a blastocyst after in vitro fertilization of MT2C_Mm KO mice. *Pcgf5^ΔMT2C_Mm/ΔMT2C_Mm^* sperm and wild-type oocytes were used for *Pcgf5^ΔMT2C_Mm/+^* embryos. Scale bar, 100 μm. (G) Development rate of embryos in each stage after in vitro fertilization of MT2C_Mm KO mice. The embryos reaching the two-cell, four-cell, morula, and blastocyst stages were counted at 24, 48, 72, and 96 hpi, respectively. The numbers of zygotes were set as 100%. Data are expressed as mean ± s.e.m. (n = 3). Dunnett’s test: *P < 0.05. (H) Representative images of PCGF5 immunostaining analysis in WT and MT2C_Mm KO embryos. Late two-cell embryos and four-cell embryos were collected at 36 and 48 hpi, respectively. The average signal intensity of WT was set as 1.0 in each stage. Bars overlaid on the plots indicate the mean. The numbers of embryos examined (n) are indicated. Dunnett’s test: *P < 0.05, ***P < 0.001; n.s., not significant. Scale bar, 20 μm. (I) Representative images of H2AK119ub1 and H3K27me3 immunostaining analysis in WT and MT2C_Mm KO embryos. Late two-cell embryos and four-cell embryos were collected at 36 and 48 hpi, respectively. The average signal intensity of WT was set as 1.0 in each stage. Bars overlaid on the plots indicate the mean. The numbers of embryos examined (n) are indicated. Dunnett’s test: ***P < 0.001; n.s., not significant. Scale bar, 20 μm. (J) Expression changes in each *Pcgf5* variant in MT2C_Mm KO embryos. Data were obtained using RT-qPCR, and the value of wild-type embryos was set to 1. Bars show means with standard error of the mean (s.e.m.). Biological replicates (n) are indicated. Dunnett’s test : *P < 0.05; U.D., undetected.

### *Pcgf5* knockdown experiments using ASOs

In *Pcgf5* KD experiment using ASOs, KD embryos exhibited embryonic lethality from the two-cell to four-cell stage and these KD two-cell embryos had a more significant decrease in the levels of both total *Pcgf5* and *Pcgf5^MT2C_Mm^* transcripts than siRNA-mediated *Pcgf5* KD embryos at the late two-cell stage (Fig. 7A–D). Furthermore, PCGF5, H2AK119ub1, and H3K27me3 protein levels were also suppressed up to 72 hpi, when embryonic development was stopped (Fig. 7E–G). Interestingly, several genes expressed during ZGA were upregulated, including MERVL (Fig. 7E).

**Fig. 7.**
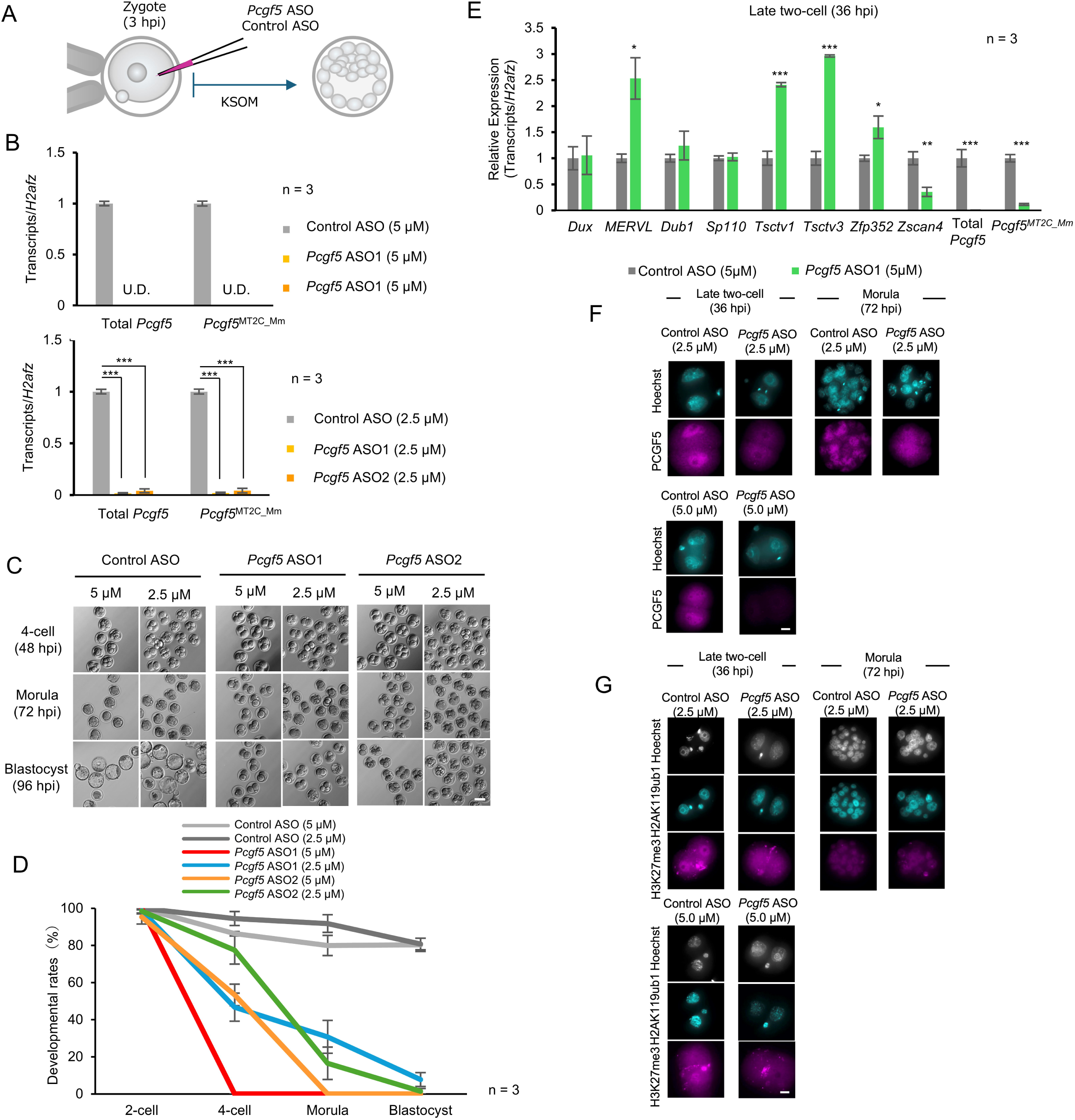
Analysis of ASO-mediated *Pcgf5* KD embryos. (A) Scheme of zygote microinjection for targeting zygotic *Pcgf5* mRNA with ASO. (B) RT-qPCR analysis of total *Pcgf5* and *Pcgf5*^MT2C_Mm^ mRNA levels in late two-cell embryos microinjected with Control ASO, *Pcgf5* ASO-1, or *Pcgf5* ASO-2 at 3 hpi (2.5 µM and 5 µM). The amount of transcripts in Control ASO-microinjected embryos was defined as 1, and the expression levels were normalized to *H2afz* as an internal control. Data are expressed as mean ± s.e.m. (n = 3). Dunnett’s test: ***P < 0.001; U.D., undetected. (C) Phase-contrast images for control ASO-microinjected (both 2.5 and 5 μM) and *Pcgf5* ASO-microinjected (2.5 μM and 5 μM) embryos after 48, 72, and 96 hpi. Scale bar, 100 μm. (D) Survival of embryos per preimplantation stage 96 h after microinjection from (B). (E) Genes expressed during ZGA as measured by RT-qPCR in Control ASO and *Pcgf5* ASO KD late two-cell embryos at 36 hpi. Many major genes were significantly upregulated in *Pcgf5* ASO KD embryos. Two-tailed Student’s t-test: *P < 0.05, ***P < 0.001. (F) Representative images of PCGF5 immunostaining in CTR and *Pcgf5* ASO-1 (2.5 µM and 5 µM)-microinjected embryos. Late two-cell embryos, four-cell embryos, and morulae were collected at 36, 48, and 72 hpi, respectively. Scale bar, 20 μm. (G) Representative images of H2AK119ub1 and H3K27me3 co-immunostaining in CTR and *Pcgf5* ASO-1 (2.5 µM and 5 µM)-microinjected embryos. Late two-cell embryos, four-cell embryos, and morulae were collected at 36, 48, and 72 hpi, respectively. Scale bar, 20 μm.

## Discussion

The total *Pcgf5* transcript expression results at each stage obtained in this study (Fig. 1B) were concordant with the RNA-seq data from a previous study (Fig. S1). Almost all *Pcgf5* transcripts were *Pcgf5^MT2C_Mm^* in the late two-cell stage, whereas variants other than *Pcgf5^MT2C_Mm^* were more highly expressed in the blastocyst stage (Fig. 1C), indicating that the transcriptional mechanism of *Pcgf5* changes dramatically during early embryogenesis. These data suggest that MERVL chimeric *Pcgf5* contributes to the function of *Pcgf5* in late two-cell stage embryos.

Previous study showed that one spot of *Xist* RNA per blastomere was detected only in female embryos at the morula stage,^16^ as in Fig. 2D. The fact that PCGF5 accumulates closest to the inactivated X chromosome in mouse embryonic stem cells^24^ and our observation that the immunofluorescence images of PCGF5 protein and *Xist* staining are similar in morulae (Fig. 2C, D) suggest that PCGF5 may also contribute to XCI by accumulating in inactivated X chromosome during the morula stage. An elevation in *Xist* expression and a reduction in the H3K27me3 modification level in the maternal *Xist* allele in *Pcgf5* siRNA KD embryos (Figure 5C, D) suggest that *Pcgf5* plays a role in maintaining non-canonical imprinting on the maternal *Xist* allele and is involved in XCI during mouse embryogenesis. This study is the first report that XCI is regulated by retrotransposons, as *Pcgf5* expression is induced by the MERVL promoter. Because few *Pcgf5* ASO KD embryos developed into morulae, non-canonical imprinting gene analysis could not be performed.

We revealed that decreased expression of *Pcgf5* led to increased expression of the major ZGA genes at the four-cell stage (Fig. 4B). A previous study revealed that oocytes lacking ubiquitin-specific peptidase 16 (Usp16), the major deubiquitinating enzyme for H2AK119ub1 in mouse oocytes, have defects in undergoing ZGA or gaining developmental competence after fertilization, despite conditional KO of Usp16 in oocytes having little effect on their survival, growth, or meiotic maturation,^28^ and these findings suggest that H2AK119ub1 is involved in the progression of ZGA. In our study, decreased H2AK119ub1 level was also observed in *Pcgf5* KD morulae, which may affect the upregulation of ZGA genes (Fig. 5A, B). Because *Pcgf5* is one of the MERVL-driven ZGA genes, *Pcgf5* expression may itself also be regulated by H2AK119ub1, and this negative feedback mechanism might lead to the short-term expression of ZGA genes. In addition, a previous study showed that maternal *Eed* KO embryos successfully developed to the blastocyst stage, even though the noncanonical imprinting genes (*Gab1*, *Phf17/Jade1*, *Sfmbt2*, *Smoc1*, and *Slc38a4*) were abnormally expressed.^29^ The significant decrease in the developmental rate in *Pcgf5* siRNA KD embryos in our study might be due to the abnormal expression of ZGA genes rather than that of noncanonical imprinting genes.

It was recently reported that ASOs, in contrast to siRNA, can degrade nuclear MERVL transcripts and cause embryonic lethality.^8^ As the developmental arrest occurred earlier in the ASO experiments compared to the siRNA experiments in our results (Figures 3B, C, 7C, D), suggesting that the *Pcgf5* transcript prior to translation may play a crucial role in the nucleus, and/or that the degradation of nuclear RNA by ASO may more efficiently suppress the *Pcgf5* transcript. Additionally, suppression of PCGF5 protein by siRNA KD was found only during a short period, compared to ASO mediated KD continued up to 72 hpi (Fig. 3D, E, 7F), and it may be because the expression of *Dicer1*, which is essential for siRNA function, is decreased after the two-cell stage in mouse embryos.^30^ As all *Pcgf5* ASO KD embryos ceased development at the ZGA stage and no *Pcgf5* homozygous KO mice were born at all (Figs. 6B and 7B, E), it suggests that if *Pcgf5* is completely suppressed, development will not proceed to morulae and that problems may occur in the ZGA stage.

The *Pcgf5^ΔMT2C_Mm/ΔMT2C_Mm^* mice generated in this study did not show a severe phenotype compared with *Pcgf5^ΔEx^*^5^*^/ΔEx^*^5^ or *Pcgf5* ASO KD embryos (Fig. 6B), which seems to be due to the complementary expression of *Pcgf5^CAN^*in *Pcgf5^ΔMT2C_Mm/ΔMT2C_Mm^* embryos (Fig. 6H). Embryos from *Pcgf5^ΔMT2C_Mm/ΔMT2C_Mm^* mice, expressed half of total *Pcgf5* mRNA (Fig. 6J), suggesting that, when some amount of PCGF5 remains in the two-cell stage embryo, these embryos could develop to the morula stage (Fig. S3). If these *Pcgf5*-reduced embryos reach the morula stage, XCI may be affected because the *Pcgf5^ΔMT2C_Mm/ΔMT2C_Mm^* and *Pcgf5^ΔEx^*^5^*^/+^* mice were less likely to born females (Fig. 6D, E), which is consistent with XCI deficient mice phenotype.^24^

In conclusion, *Pcgf5* functions to regulate genes expressed during ZGA via H2AK119ub1 and H3K27me3 modifications, and its defective modifications may be a contributing factor to developmental delay. *Pcgf5* was also found to have a role in maintaining the H3K27me3 modification on the maternal *Xist* region.

### Experimental model and subject details In vitro fertilization and embryo culture

Slc:ICR (Japan SLC, Hamamatsu, Japan) mice were used for all experiments except for *Pcgf5* KO experiments and ChIP-seq analysis. Seven- to 10-week-old female mice were superovulated by injection of 7.5 IU of equine chorionic gonadotropin (ASKA Pharmaceutical, Tokyo, Japan) followed by 7.5 IU of human chorionic gonadotropin (ASKA Pharmaceutical) 48 h later. Cumulus oocyte complexes were harvested 14 h after human chorionic gonadotropin injection and placed in human tubal fluid medium supplemented with 4 mg/ml bovine serum albumin (BSA, A3311; Sigma-Aldrich, St. Louis, MO) covered with paraffin oil (Nacalai Tesque, Kyoto, Japan). Spermatozoa were collected from the cauda epididymis of 9–16-week-old male mice and cultured in human tubal fluid medium for 1 h. After capacitation, sperm were introduced into oocyte-containing droplets at a final concentration of 1 × 10^6^ cells/ml. After a 3-h incubation at 37°C in a 5% CO_2_ atmosphere, fertilized oocytes were washed with K^+^-modified simplex optimized medium (KSOM) supplemented with amino acids^31^ and 1 mg/ml BSA and then used for microinjection or cultured in the same medium under paraffin oil at 37°C in a 5% CO_2_ atmosphere to the following stages: one-cell (12 h hpi), early two-cell (24 hpi), late two-cell (36 hpi), four-cell (48 hpi), eight-cell (54 hpi), morula (72 hpi), and blastocyst (96 hpi). MII oocytes were collected from cumulus oocyte complexes followed by treatment with 1% hyaluronidase to remove cumulus cells.

### Collection of mouse preimplantation embryos for ChIP-seq analysis

For superovulation, 4- to 10-week-old C57BL/6JmsSlc (Japan SLC) females were injected with 0.15 ml CARD HyperOva followed by 5 IU of human chorionic gonadotropin (ASKA Pharmaceutical Co., Tokyo, Japan) 48 h later. Spermatozoa were obtained from the cauda epididymis of adult male PWK/PhJ mice [RBRC00213 (RIKEN BioResource Research Center, Tsukuba, Japan) originated from 003715 (The Jackson Laboratory, Bar Harbor, ME)]. Subsequent steps were performed using the method described above.

### Electroporation

To perform the electroporation of zygotes, we employed the NEPA21 electroporator system (Nepa Gene Co., Chiba, Japan) with a glass slide and two metal electrodes separated by a 1-mm gap (CUY501P1-1.5, Nepa Gene Co.). The electroporation parameters consisted of four poring pulses (40 V; pulse length, 1.0, 2.0, 3.0, or 4.0; interval, 50 msec; decay rate, 10%; polarity, +) and five transfer pulses (5 V; pulse length, 50 msec; interval, 50 msec; decay rate, 40%; polarity, +/−). Opti-MEM (Thermo Fisher Scientific, Waltham, MA, USA) containing si*Pcgf5^CAN^*-1, si*Pcgf5^CAN^*-2, or siControl (siRNAs at 5 µM each; RNAi Inc., Tokyo, Japan) was prepared as the siRNA solution. The prepared siRNA solution (4.5 µl) was applied between the two electrodes on a glass slide. After being washed with 50 µl Opti-MEM, zygotes were aligned between the electrodes and an electrical discharge was applied. The siRNA solution was exchanged every two operations to avoid dilution. After electroporation, the zygotes were transferred to 50 µl Opti-MEM and washed in KSOM. The siRNA sequences are listed in Table S1.

### Plasmid construction, RNA extraction, and RT-qPCR

Based on the transcription start site of *Pcgf5^CAN^* predicted from the NCBI database (NM_001368690.1, NM_029508.4, NM_001360691.1, and XM_036161722.1) and that of *Pcgf5^MT2C_Mm^*predicted from previously reported RNA-seq data,^1^ the sequences of *Pcgf5^CAN^* and *Pcgf5^MT2C_Mm^* were cloned by PCR from the cDNA of late two-cell embryos. These fragments were incorporated into a pTAC-1 vector plasmid (TA PCR Cloning Kit, DS125; BioDynamics Laboratory, Tokyo, Japan). *H2afz* and *Gapdh* PCR fragments were cloned using the same method. The primer sequences are listed in Table S2.

RNA extraction and reverse transcription were performed using the SuperPrep II Cell Lysis & RT Kit for qPCR (Toyobo, Osaka, Japan). Briefly, 10 embryos or oocytes were washed three times in PBS containing 0.5% polyvinylpyrrolidone (PVP K30; Nacalai Tesque) (PVP/PBS) and collected in PCR tubes along with 0.5 µl of PVP/PBS. After adding 3.5 µl of lysate, all lysates were subjected to reverse transcription. Transcription levels were measured using the StepOnePlus real-time PCR system (Thermo Fisher Scientific) with KOD SYBR qPCR Mix (Toyobo), and relative gene expression was calculated using the 2^−ΔΔCt^ method normalized to the corresponding *H2afz* levelz.^32,33^ Absolute expression was determined using serial dilutions of the plasmids containing the cloned chimeric *Pcgf5* and *Pcgf5* variants for in vitro transcription or using the pTAC plasmid in which an *H2afz* and *Gapdh* PCR fragment was cloned. The primers used for quantification are listed in Table S2.

### Immunofluorescence

For PCGF5 immunofluorescence, embryos were treated with acid Tyrode’s solution (800 mg NaCl, 20 mg KCl, 15 mg CaCl_2_, 10 mg MgCl_2_•6H_2_O, 4 mg Na_2_HPO_4_, 100 mg Glucose, 400 mg PVP K30, 3.62 ml 5N HCl, up to 100 ml water) to remove their zona pellucida. These embryos were then fixed in PBS containing 4% paraformaldehyde for 20 min at 28°C. After three washes in PVP/PBS, the embryos were permeabilized with 0.5% Triton X-100 (Sigma-Aldrich) in PBS for 40 min at 28°C. After washing three times in PVP/PBS, the embryos were blocked in PBS containing 1.5% BSA, 0.02% Tween 20, and 0.2% sodium azide (blocking buffer) for 1 h at 28°C and then incubated overnight at 4°C with a mouse anti-PCGF5 antibody (1:250 dilution, ab201511; Abcam, Cambridge, UK). After three washes in blocking buffer, the embryos were incubated in blocking buffer containing Alexa Fluor 555 donkey anti-rabbit IgG secondary antibody (1:500 dilution, A31572; Thermo Fisher Scientific) for 1 h at 28°C. Immunostained embryos were washed three times in blocking buffer, and nuclei were stained in blocking buffer containing 1 mg/ml Hoechst 33342 (Merck KGaA, Darmstadt, Germany,) for 10 min at 28°C. After a 5-min wash with blocking solution, the embryos were transferred and mounted by dropping 2.5 µl of VECTASHIELD (Vector Laboratories, Newark, CA) onto a slide glass and covered with cover glass. A fluorescence microscope (IX73; Olympus Inc., Tokyo, Japan) was used to observe fluorescence signals.

For H2AK119ub1 and H3K27me3, embryos were treated in acidic Tyrode’s solution to remove their zona pellucida, fixed in 3.7% paraformaldehyde in PBS containing 0.2% Triton X-100 at 28°C, and blocked in blocking buffer for 1 h at 28°C. After three washes with blocking buffer, the embryos were incubated overnight at 4°C in blocking buffer containing rabbit anti-H2AK119ub1 antibody (1:2,000 dilution, 8240; Cell Signaling Technology, Danvers, MA) and mouse anti-H3K27me3 antibody (1:500 dilution, 61017; Active Motif, Carlsbad, CA). After washing three times in blocking buffer, the embryos were incubated in blocking buffer containing secondary antibodies (Alexa Fluor™ Plus 488 goat anti-mouse IgG, 1:500 dilution, A32723, Thermo Fisher Scientific; and Alexa Fluor 555 donkey anti-rabbit IgG, 1:500 dilution, A31572, Thermo Fisher Scientific) for 1 h at 28°C. Subsequent Hoechst staining was performed as described above.

### Xist probe for FISH

A probe for *Xist* RNA was prepared by using Nick Translation DNA Labeling System 2.0 (ENZ-GEN111-0050; Enzo, Long Island, NY) with Red 580 dUTP (ENZ-42844; Enzo) or Green 496 dUTP (ENZ-42831; Enzo). The template DNA was pCMV-Xist-PA (26760; Addgene, Watertown, MA). pCMV-Xist-PA was a gift from Rudolf Jaenisch (Addgene plasmid # 26760 ; http://n2t.net/addgene:26760 ; RRID:Addgene_26760).^34^ The fluorescent probes were ethanol-precipitated with 5 µg of mouse Cot-1 DNA (2402609; Invitrogen, Carlsbad, CA), 5 µg of herring sperm DNA (0000538816; Invitrogen), and 2.5 µg of yeast tRNA (01333800; Invitrogen) and then dissolved in 20 µl of formamide (M2N0838; Nacalai Tesque). The probes were stored at 4°C. Before being used, the probes (0.75 µl each) were mixed with 0.75 µl of Cot-1 DNA/formamide and 2.25 µl of 4× SSC/20% dextran (3905320; Merck KGaA). The probe mixtures were heated for 30 min at 80°C and then transferred to a 37°C incubator (“preannealed probes”).

### Whole-mount RNA FISH

Whole-mount RNA FISH was basically performed as previously described. Cover glasses used for FISH were soaked in 50× Denhardt’s solution (10727-74; Nacalai Tesque) diluted with nuclease-free water (1039498; Qiagen, Hilden, Germany) and incubated at 65°C for 3 h. These cover glasses were then placed in a 3:1 mixture of methanol (Fujifilm Wako Pure Chemical Corp., Osaka, Japan) and glacial acetic acid (Fujifilm Wako Pure Chemical Corp.) for 20 min at room temperature. The solution was then air-dried and stored at 4°C.

Morulae were fixed at 78 hpi in acidic Tyrode’s solution to remove their zona pellucida. The samples were then washed three times with PVP/PBS and placed on Denhardt’s-treated cover glass with PVP/PBS using a glass pipette, and the surrounding PVP/PBS was air-dried by aspirating the PVP/PBS with a glass pipette. Next, 4% paraformaldehyde in PBS was dropped onto the samples and left for 20 min at room temperature. After three washes with 1× PBS, the embryos were treated with 0.1 N HCl containing 0.02% Triton X-100 for 15 min at 4°C. After three washes with 1× PBS, the embryos were then covered with 4.5 µl of “preannealed probes” and incubated for > 24 h at 37°C. The embryos were washed three times with 0.1% BSA/2× SSC prewarmed to 42°C for 15 min each and twice with 2× SSC prewarmed to 42°C for 15 min each and then mounted on a glass slide in VECTASHIELD® PLUS Antifade Mounting Medium with DAPI (H-2000; Vector Laboratories). A fluorescence microscope (BX73; Olympus Inc.) was used to observe fluorescence signals.

### Assessment of the mouse embryo development rate

Embryos were incubated for 96 h after insemination, and the embryo development percentage was evaluated by observing the embryos under a microscope every 24 h. Normal embryos were defined as those that had formed two blastomeres at 24 h, four blastomeres at 48 h, compaction at 72 h, and a blastocoel at 96 h after insemination.

### Low-input ChIP-seq library preparation

At 78 hpi, ∼40 morulae per group were briefly treated with acidic Tyrode’s solution to remove the zona pellucida, washed with PVP/PBS, and transferred to 1.7-ml tubes. Basically, the ChIP was performed using H3K27me3 antibody (C15410069; Diagenode, Seraing, Belgium) and as previously described.^35^ The index was added to the samples based on the protocol of the ThruPLEX DNA-Seq Kit (Takara Bio Inc., Kusatsu, Japan). The amplification was performed for 18 cycles.

### Sequencing and data analysis

After library preparation, samples were sequenced on a NovaSeq X (Illumina, Inc., San Diego, CA). These reads were quality-checked and adapter-trimmed using fastp (ver. 0.20.1).^36^ The reads were then mapped to the N-masked C57BL/6J mouse genome (mm10) using single nucleotide polymorphisms (SNPs) with PWK mice (https://www.ebi.ac.uk/pub/databases/eva/PRJEB43298/mgp_REL2005_snps_indels.vcf.gz) processed through bowtie2 (ver. 2.4.5) with “-t-q -N 1 -L 25 --no-mixed --no-discordant” options.^22,37^ The reads of the generated BAM files were sorted to C57BL/6J or PWK mice using UniversalSNPsplit (https://github.com/lanjiangboston/). BAM files comprising reads specific to each strain were converted to bigwig (BW) files using counts per million (CPM) normalization by bamCoverage in deepTools.^38^ ChIP sample peaks were obtained by MACS2 (ver. 2.2.7.1) by comparison to baseline.^39^ Landscapes of gene-specific ChIP peaks were visualized by loading BW files into Integrative Genomics Viewer (IGV) (ver. 2.12.1).

### Generation of *Pcgf5* and *Pcgf5^MT2C_Mm^* KO mice

All KO mouse strains were generated in a C57BL/6JmsSlc (Japan SLC) background. To generate *Pcgf5* KO mice, two gRNAs designed to intercept exon 5 (Table S3), which is common to all *Pcgf5* variants, and Cas9 protein (Cas9 nuclease protein NLS, 319-08641: Nippon Gene, Tokyo, Japan) were electroporated into 3 hpi embryos, and two-cell embryos were transferred into the oviduct of pseudo-pregnant ICR females.^40^ *Pcgf5^MT2C_Mm^* KO mice were similarly generated by electroporation using gRNAs that intercepted only the MT2C_Mm sequence (Table S3). KO mice were confirmed through genotyping and Sanger sequencing (Table S4). The primer sequence used for Sanger sequencing is as follows: Sanger_1_F, 5′-TAAGTTGGGTAACGCCAGGG -3′. The obtained KO mice were crossed once with wild-type C57BL/6J mice, and the acquired heterozygous KO mice were crossed with each other to produce the homogenous KO mice.

### Design of *Pcgf5*-targeting ASOs

Two independent 3-10-3 gapmer ASOs were designed based on the consensus sequence of *Pcgf5* (Table S3 and S5) and ordered from Integrated DNA Technologies (Coralville, IA). The full-length *Pcgf5* mRNA sequences were entered into Sfold’s Soligo (https://sfold.wadsworth.org/cgi-bin/soligo.pl), which selected sequences with GC percentages of 40%–60%, a binding energy of less than −8 kcal/mol, and no GGGG sequences.^41^ RNAfold (http://rna.tbi.univie.ac.at/cgi-bin/RNAWebSuite/RNAfold.cgi) was then used to predict the secondary structure of the RNA to see if the target sequences were on a single-stranded loop of the full-length *Pcgf5* mRNA.^42^ Finally, non-specific binding was checked using NCBI Blast (https://blast.ncbi.nlm.nih.gov/Blast.cgi) to determine whether the candidate sequences bound to mRNA and genomic DNA besides *Pcgf5*.

## Microinjection of ASOs

The above-designed ASOs were diluted in water at 2.5 µM or 5 µM concentration and microinjected at approximately 3–5 pL into the cytoplasm of zygotes at 3 hpi in HEPES-buffered KSOM. Microinjection was performed under an inverted microscope (IX73) equipped with a piezo injector (PMAS-CT150; PrimeTech, Tokyo, Japan), FemtoJet 4i (Eppendorf, Hamburg, Germany) and a micromanipulator (IM-11-2; Narishige, Tokyo, Japan).

### Statistical analyses

Each experiment was repeated at least three times. Quantitative and absolute RT-qPCR and quantification of immunofluorescence data were analyzed using a Student’s *t*-test for pairwise comparisons. All analyses were performed using R (ver. 4.2.1), and significance was accepted at P < 0.05. All fluorescence intensity analyses were performed with ImageJ.

## Ethical approval for the use of animals

All animal experiments were approved by the Animal Research Committee of Kyoto University (Permit Numbers: R4-17 and R5-17) and were performed in accordance with the committee’s guidelines.

## Resource availability Lead contact

Further information and requests for resources and reagents should be directed to and will be fulfilled by the lead contact, Shinnosuke Honda.

## Materials availability

This study did not generate any new unique reagents.

## Data and code availability

The data of this study is available from the corresponding author upon request. NGS data reported in this study have been deposited at Gene Expression Omnibus (GEO) under accession no. GSE280946. This paper does not report the original code.

## Acknowledgements

This work was supported by a Grant-in-Aid for Research Activity Start-up (no.22K20612) from the Japan Society for the Promotion of Science, and a grant from the Livestock Promotional Subsidy from the Japan Racing Association.

## Author contributions

Conceptualization, S.M., N.M. and S.H.; methodology, S.M., S.I. and S.H.; software, S.M.; validation, S.M., T.Y.; formal analysis, S.M.; investigation, S.M.; data curation, S.M.; writing – original draft, S.M.; writing – review & editing, S.M., T.Y., N.M., S.I. and S.H.; visualization, S.M.; supervision, S.H.; project administration, S.I. and S.H.; funding acquisition, S.I. and S.H.

