## Supplemental file for "Endogenous retrovirus-driven *Pcgf5* plays critical roles in zygotic genome activation and noncanonical imprinting"

**Supplementary Data**

Supplementary Fig. 1

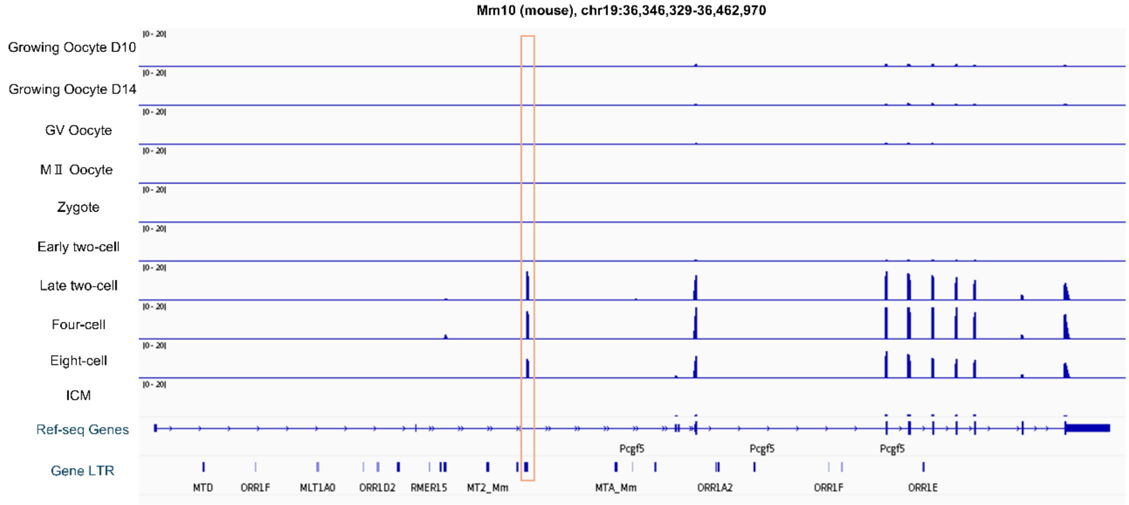


Genome browser view of RNA-seq data from each stage of mouse preimplantation embryos. Data are presented on growing oocytes day 10 (D10), growing oocytes day 14 (D14), GV oocytes, MII oocytes, zygotes, early two-cell embryos, late two-cell embryos, four-cell embryos, eight-cell embryos, and the inner cell mass of blastocysts. The *Pcgf5*$^{CAN}$ exon and LTR retrotransposon locations are indicated at the bottom. MT2C_Mm for the chimeric expression of *Pcgf5*$^{MT2C\_Mm}$ is shown as an orange square.

Supplementary Fig. 2

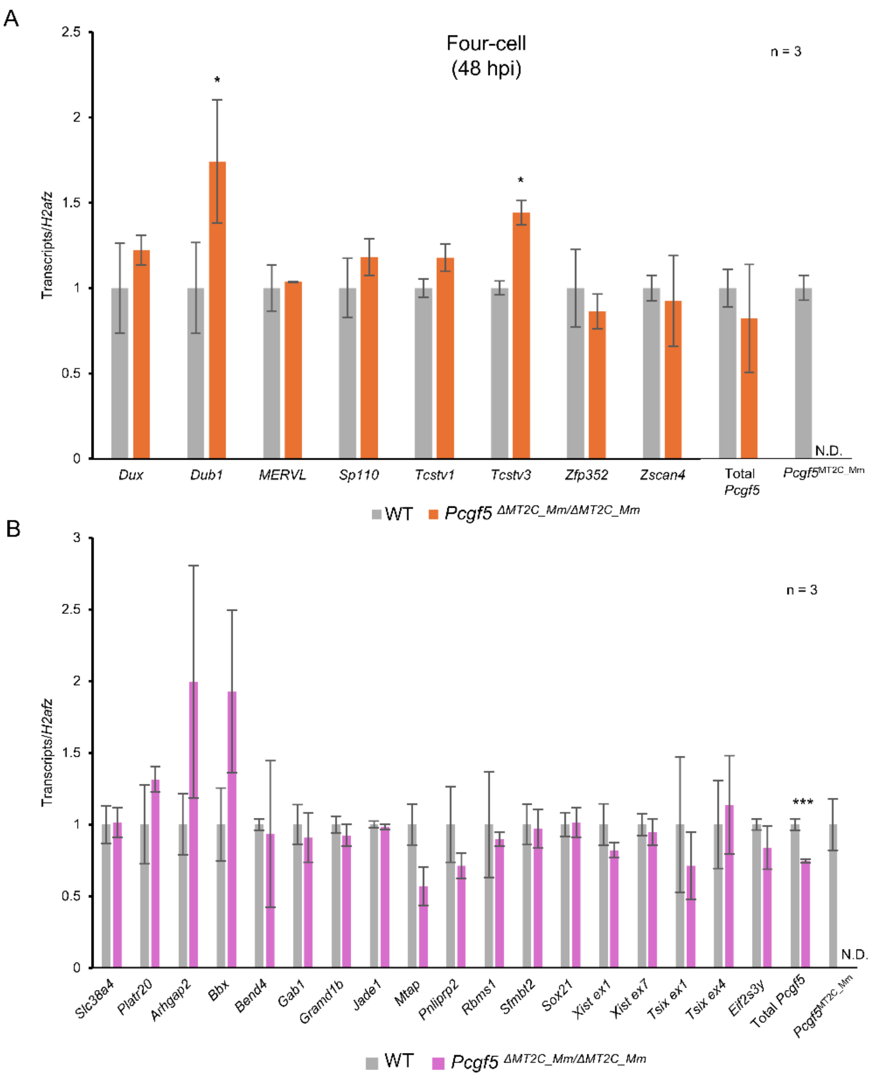


(A) Genes expressed during ZGA as measured by RT-qPCR in wild-type and MT2C_Mm homozygous KO late two-cell embryos at 36 hpi. Two-tailed Student's t-test: *P < 0.05; N.D., not detected.

(B) Maternal noncanonical imprinting genes as measured by RT-qPCR in wild-type and MT2C_Mm homozygous KO morulae at 72 hpi. Two-tailed Student's t-test: *P < 0.05, ***P < 0.001; N.D., not detected.

Supplementary Fig. 3

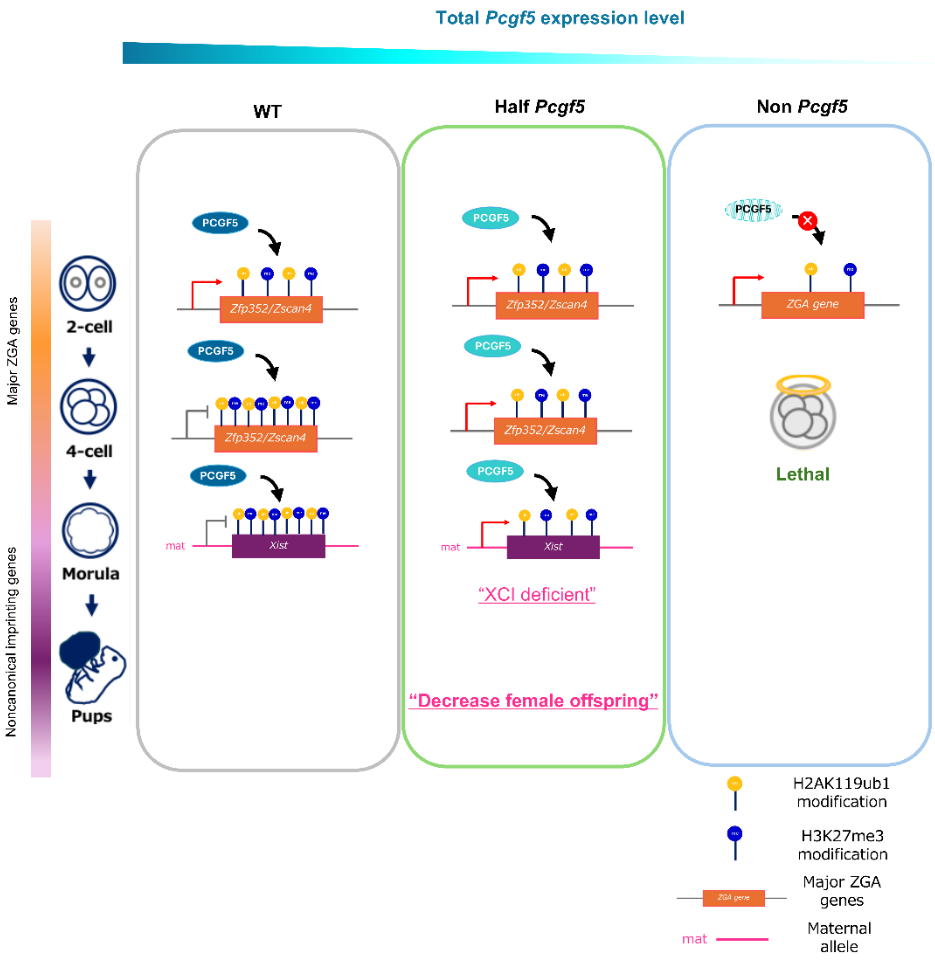


**Dynamic models and functions of PCGF5 in mouse preimplantation embryos.**

These models illustrate the overall *Pcgf5* dynamics from the two-cell stage to the morula stage and the predicted mechanisms of regulation of ZGA genes and H3K27me3 imprinting genes for all of the *Pcgf5* variants, including chimeric *Pcgf5*. Which of the two phenotypes dominates depends on the amount of total *Pcgf5* mRNA in the two-cell stage.

Supplementary Table 1. Sequences of siRNAs used for electroporation

| Names of siRNAs | Sense Strand (5′→3′) | Antisense Strand (5′→3′) |
|---|---|---|
| *siPcgf5*$^{can}$-1 | GGAGAAAAUUCAUACCCGAtt | UCGGGUAUGAAUUUUCUCCtt |
| *siPcgf5*$^{can}$-2 | GGUCUCAUGAAGAAAUUUAtt | UAAAUUUCUUCAUGAGACCtt |
| siControl | UUCUCCGAACGUGUCACGUtt | ACGUGACACGUUCGGAGAAtt |

Supplementary Table 2. Primer sequences for quantitative PCR

| Primer names | | Sequences | Product sizes (bp) |
|---|---|---|---|
| *Total Pcgf5* | sense | CTGATCAAGCCCACGACAGT | 110 |
| | anti-sense | TGAACTTGGTTGCCACACCT | |
| *Pcgf5*$^{MT2C\_Mm}$ | sense | GGGACTGAACAACCGCTAGA | 148 |
| | anti-sense | TTCCTGTAGGAGTCCCGAGG | |
| *NM90* | sense | CCTCCCGAGAGCTAGGTATGAG | 105 |
| | anti-sense | TTCCTGTAGGAGTCCCGAGG | |
| *NM08* | sense | CCATCCCATCGTGCATCTCA | 138 |
| | anti-sense | TTCCTGTAGGAGTCCCGAGG | |
| *NM91* | sense | TTTCCCGGGCTCGGAGG | 147 |
| | anti-sense | TTCCTGTAGGAGTCCCGAGG | |
| *XM22* | sense | TTGGGATTGGGACTGCGTTT | 140 |
| | anti-sense | TTCCTGTAGGAGTCCCGAGG | |
| *Dux* | sense | GGCCCTGCTATCAACTTTCAAGA | 135 |
| | anti-sense | TCCTCTCCACTGCGATTCCT | |
| *Dub1* | sense | GGAGACATGGTGGTTGCTCT | 140 |
| | anti-sense | CTCTCCCAACTCAGACTGTGC | |
| *Sp110* | sense | AAGGATCCAGGAACCCCTTA | 159 |
| | anti-sense | GCATAGGCGATGTTCACCTT | |
| *Tcstv1* | sense | TGAACCCTGATGCCTGCTAAGACT | 102 |
| | anti-sense | AGATGGCTGCAAAGACACAACTGC | |

| | | | |
|---|---|---|---|
| *Tcstv3* | sense | AGAAAGGGCTGGAACTTGTGACCT | 109 |
| | anti-sense | AAAGCTCTTTGAAGCCATGCCCAG | |
| *MERVL* | sense | CTCTACCACTTGGACCATATGAC | 83 |
| | anti-sense | GAGGCTCCAAACAGCATATCTA | |
| *Zscan4* | sense | GAGATTCATGGAGAGTCTGACTGATGAGTG | 134 |
| | anti-sense | GCTGTTGTTTCAAAAGCTTGATGACTTC | |
| *Zfp352* | sense | AAAGCCTTGATCCTCAGGTG | 115 |
| | anti-sense | GCCGAAGAGTTTTTCTGAGG | |
| *Msfd7c* | sense | GTCCTTGCTTGGTCTCTTGC | 168 |
| | anti-sense | CTTCCTCTCGTGACCCTCAG | |
| *Slc38a4* | sense | GGCTTCTTCTGCCACTATGC | 108 |
| | anti-sense | AAGACCAAAGCCCCAATCTT | |
| *Platr20* | sense | CTGCACGCAAGTCTGTTCAA | 102 |
| | anti-sense | CACTCTGCTTTCCCGTCTGG | |
| *Arhgap2* | sense | TGCCTGCTGAGTCTACTGCTA | 147 |
| | anti-sense | TCTGTTTTGTTGGGCGAGACT | |
| *Bbx* | sense | GTGTCACAACCAGCTCAGTAGAC | 145 |
| | anti-sense | CTGCCTTTCATTACTGTGACCAG | |
| *Bend4* | sense | TCAGGAGAGACAGGTCCCG | 145 |
| | anti-sense | CAGCATCGCTGAACACTTTGT | |
| *Gab1* | sense | TTCAGGTCCAGCCCAAAGAC | 147 |
| | anti-sense | AGTTCGCTCTAAACCATGGGG | |
| *Gramd1d* | sense | GCTGCTGGTTATCAGCTGTG | 148 |
| | anti-sense | GGGTAACCTTTCCTGGAGCC | |

| | | | |
|---|---|---|---|
| *Jade1* | sense | CCCTGAGGTAAGCATTGGCA | 147 |
| | anti-sense | GTCCGGCAGTTCTTCACAGA | |
| *Mtap* | sense | GAGATCCAGCCTGGTGACAT | 142 |
| | anti-sense | TTTTGGGGCAAAACGGTTCA | |
| *Pnliprp2* | sense | AGGTCGGCCATCTGGATTTC | 123 |
| | anti-sense | AGGCTGCGAAGTTTCGAGTT | |
| *Rbms1* | sense | CTTGGTTACCTTTGGCCTGCTG | 131 |
| | anti-sense | CAGCTGATCCCATCCTGAGTTG | |
| *Sfmbt2* | sense | TTGACTGGTTCTCGGACAGC | 110 |
| | anti-sense | TGGGAGTCCTGCAGCTCTAT | |
| *Sox21* | sense | GCCGGTGACTCGTGTCTTTA | 168 |
| | anti-sense | GAACGGCGGTCATCTCTCAT | |
| *Xist ex1* | sense | GCCAACCAATGAGACCACTT | 129 |
| | anti-sense | TTCTCTCAAACCACCACACG | |
| *Xist ex7* | sense | CTTTGGGCTTAGGTGAGCAG | 123 |
| | anti-sense | CCAGGAGTTCCTTTGGTGAA | |
| *Tsix ex1* | sense | TACCTGCAAGCGCTACACAC | 142 |
| | anti-sense | GCTGGCTATCACGCTCTTCT | |
| *Tsix ex4* | sense | CGACCTCAGATGAGGAGAGG | 96 |
| | anti-sense | CCACCAAATCGGTCACAACT | |
| *Eif2s3y* | sense | AATTGCCAGGTTATTTTCATTTTC | 151 |
| | anti-sense | AGTTCAGTGGTGCACAGCAA | |
| *H2afz* | sense | GGTAAAGCGTATCACCCCTCG | 130 |
| | anti-sense | CTTCCCGATCAGCGATTTGTG | |

| | | | |
|---|---|---|---|
| *Gapdh* | sense | TGTCATCATACTTGGCAGGT | 73 |
| | anti-sense | GACACATTGGGGGTAGGAACACG | |

Supplementary Table 3. Sequences of gRNA for generating KO mice

| Names of gRNA | Sequences (5′→3′) |
|---|---|
| *Pcgf5* gRNA1 | ggtaattgaagcttcataac |
| *Pcgf5* gRNA2 | ggcacgagccaactccttca |
| MT2C_Mm gRNA1 | ctgtttgaggcattacgtga |
| MT2C_Mm gRNA2 | ttagatcgaatgtactcacg |

Supplementary Table 4. Primer sequences for KO mouse genotyping

| Primer names | | Sequences | Product sizes (bp) |
|---|---|---|---|
| Total *Pcgf5* | sense | AGTGTACCTGAAGTGTTGAAGGAAAGG | 1559 |
| genotyping | anti-sense | AAACTTGACTTGAGCGTGCTCTAGAAT | |
| MT2C_Mm | sense | AGAATCCAACCCCATGTGTAAAAGTAT | 869 |
| genotyping | anti-sense | CATAAAAGGTAAAACAGGCCCTGATAC | |

Supplementary Table 5. Sequences of ASOs used for microinjection

| Names of ASOs | Sequences (5′→3′) |
|---|---|
| *Pcgf5* ASO1 | **T*C*T***T*G*T*T*C*T*C*G*T*A***G*T*C*** |
| *Pcgf5* ASO2 | **A*C*T***G*C*C*C*A*T*T*A*T*T***T*C*G*** |
| Control ASO | **G*G*C***T*A*C*T*A*C*G*C*C*G***T*C*A*** |

* indicates a phosphorothioate bond. Bold residues represent locked nucleic acids (LNAs).
